# Poly(UG)-tailed RNAs are involved in the control of thousands of genes predominantly in the germline in *Pristionchus pacificus*

**DOI:** 10.1101/2025.09.01.673539

**Authors:** Veroni de Ree, Dorothee Harbecke, Hanh Witte, Christian Rödelsperger, Catia Igreja, Adrian Streit

## Abstract

In the nematode *Caenorhabditis elegans*, the terminal transferase RDE-3 adds a poly(UG)-tail to the free 3’ end of RNA molecules generated by the action of primary siRNAs or piRNAs. The tailed RNA serves as a template for RNA-dependent RNA polymerases (RdRp) to generate secondary siRNAs, thereby reinforcing the RNAi effect. In this iterative process, progressively shorter tailed RNAs are formed. In *C. elegans*, injection of poly(UG)-tailed single-stranded RNA (ssRNA) leads to RNAi-mediated gene silencing, thereby bypassing the need for the processing of double-stranded RNA into siRNAs. We wondered if poly(UG)-tailed ssRNAs could also be used for experimental gene knockdown in nematodes where long dsRNA-mediated RNAi does not work reliably, such as the satellite model organism *Pristionchus pacificus* or parasitic nematodes of the genus *Strongyloides*.

Here we show that injection of poly(UG)-tailed RNA leads to gene knock down in *P. pacificus* and that the injected RNA, as well as the corresponding endogenous RNA, serve as substrate for the formation of new poly(UG)-tailed RNAs. Different from *C. elegans*, in *P. pacificus,* the knockdown effect depends on the redundant activity of the three *rde-1* paralogs present in this species. We detected endogenously occurring poly(UG)-tailed RNAs derived from thousands of genes, more than half of which belong to germline-specific co-expression clusters. Mutations in *Ppa-rde-3*, lead to sterility. In contrast, in *Strongyloides* spp., we found poly(UG)-tailed RNAs to be much less abundant, if not absent. Our results show that poly(UG)-tailed RNAs are not restricted to *C. elegans* and suggest that they play an important function in the germ line in *P. pacificus*.

## 1 Introduction

Gene silencing in response to double stranded RNA (dsRNA) is believed to be a natural mechanism to control the activity of transposable elements that can be exploited for the experimental knock down of genes in various organisms (RNA interference [RNAi]) (Svoboda, 2020). Such work was pioneered in the model nematode *Caenorhabditis elegans* (Fire et al., 1998) and adapted to other systems (Svoboda, 2020), of which only nematodes are discussed here. In *C. elegans,* the experimental silencing is achieved by applying long (several hundred to a few thousand bp) double stranded RNA (dsRNA) by micro injection, soaking the worms in dsRNA solution or feeding the worms with dsRNA expressing bacteria (Ahringer, 2006; Conte et al., 2015). The long dsRNA is processed by Dicer and other factors into primary small interfering RNAs (siRNAs) (Seroussi et al., 2022). The siRNAs form an active complex together with Argonaute proteins, and recognize their target through base pairing (Liu et al., 2023; Seroussi et al., 2022; Seroussi et al., 2023). While very successful in *C. elegans* and most plant nematode parasites (Maule et al., 2011), many animal parasitic nematodes remained largely refractory to the approach of treatment with long dsRNA (Maule et al., 2011; Viney and Thompson, 2008). In some cases, application of *in vitro* synthesized siRNAs was successful (Dulovic and Streit, 2019; Misra et al., 2017).

A possibly nematode specific amplification mechanism is known to lead to the formation of secondary siRNAs in *C. elegans* (Pak and Fire, 2007) and other nematodes (Holz and Streit, 2017; Sarkies et al., 2015; Wang et al., 2011). These molecules are around 22 nt long, have a G at their 5’end (22G RNAs), and interact with members of the family of Worm specific Argonautes (WAGOs) (Seroussi et al., 2023). 22G RNAs are formed by RNA dependent RNA polymerases, without 5’ processing. Hence, they have a 5’ tri-phosphate that distinguishes them from other small RNAs like primary siRNAs, miRNAs or piRNAs (Pak and Fire, 2007). (Shukla et al., 2020), in a seminal paper, showed that the terminal transferase RDE-3 (also known as MUT-2) adds poly(UG) tails to target RNAs cut by primary or secondary siRNA complexes. These tailed RNAs serve as templates for the synthesis of new secondary siRNAs by RNA dependent RNA polymerases (RdRPs). These authors also showed that injection of poly(UG) tailed RNAs is sufficient to induce gene silencing. This silencing effect is independent of *rde-1*, which encodes an argonaute family protein and is a key component of the primary RNAi mechanism (Tabara et al., 1999).

Despite the success in *C. elegans*, RNAi is not routinely used in other nematode species. In the free-living model nematode *Pristionchus pacificus* RNAi has been very limited and restricted to particular genes (Aurilio and Srinivasan, 2015), although a protocol using lipofectamine in the injection mix has been proposed to improve the efficacy (Adams et al., 2019). Animal parasitic nematodes of the genus *Strongyloides* appear completely refractory to RNAi through long dsRNAs (Viney and Thompson, 2008) but moderate gene knock down by applying siRNAs has been reported (Dulovic and Streit, 2019). For both these taxa the presence of naturally occurring 5’-tri-phosphorylated small RNAs, putatively secondary siRNAs, has been shown (Holz and Streit, 2017). 22-23G secondary siRNAs are present in *P. pacificus*, while in *Strongyloides* spp. these small RNAs are longer (27 nucleotides) and initiate with G or A nucleotides (27GA RNAs).

In this study, we show that poly(UG) tailed RNAs can be used for experimental gene knock down in *P. pacificus* and that such RNAs are naturally formed from thousands of genes, many of which are expressed in the germ line.

## Results

### Injection of poly(UG)-tailed RNAs leads to loss of function phenotypes in *P. pacificus*

*P. pacificus* has an extensively studied mouth morphology polyphenism (Sommer, 2020). In response to environmental conditions, these worms form either a wide mouth with two teeth that allows them to kill other nematodes (eurostomatous, Eu) or adopt a narrower mouth with only one tooth characteristic of the strictly bacterivorous animals (stenostomatous, St). Under standard laboratory conditions on NGM plates, almost 100% of the hermaphrodites of the strain PS312 develop into Eu morphs. At the molecular level, expression of *eud-1* is essential for the formation of the Eu animals (Sommer, 2020). In a pilot experiment, we injected poly(UG)-tailed RNA targeting the gene *eud-1* (the RNA contained 464bp fragment of the *eud-1* transcript) at different concentrations, along with a plasmid encoding RFP under the control of the *eft-3* promoter, into the gonads of PS312 hermaphrodites. For the entire progeny of injected animals that produced any RFP-expressing progeny, we determined the mouth morphology (Suppl Fig. 1). In the non-injected controls, all progeny were Eu. At low poly(UG)-tailed RNA concentrations (0.17 and 0.5 pmol/µl) majority of mothers produced progeny that were only Eu (0% St). At 1.5 and 4.5 pmol/µl concentrations of poly(UG)-tailed RNA, however, the majority of injected mothers produced various numbers (4-40%) of St progeny, indicating that reduction of *eud-1* expression occurred at different levels in the new generation of worms. Since the variation of the percentage of St progeny from RFP positive injected mothers was smaller at 4.5 pmol/µl, we decided to continue our experiments with a concentration of 4.5 pmol/µl of RNA in the injection mix. This is considerably higher than the 0.5 pmol/µl used by (Shukla et al., 2020) for *C. elegans*.

In addition to *eud-1* we targeted *dpy-1,* another gene with a visually recognizable mutant phenotype (Dpy, short and fat body shape) (Witte et al., 2015). We injected poly(UG)-tailed, poly(AC)-tailed and non-tailed RNAs targeting the two genes into PS312 hermaphrodites and scored for the St and the Dpy phenotypes (Fig. 1). In both cases, we observed gene silencing as multiple St and Dpy animals were registered in the progeny of the injected animals. Importantly, in the progeny of *eud-1* injected animals we did not observe any Dpy animals and in the progeny of *dpy-1* injected animals no St worms were present, demonstrating the sequence specificity of the knock down. Strikingly, and different from the finding of (Shukla et al., 2020), RNAs without tail and RNAs with a poly(AC)-tail also induced gene silencing, although with clearly lower penetrance.

**Fig. 1:**
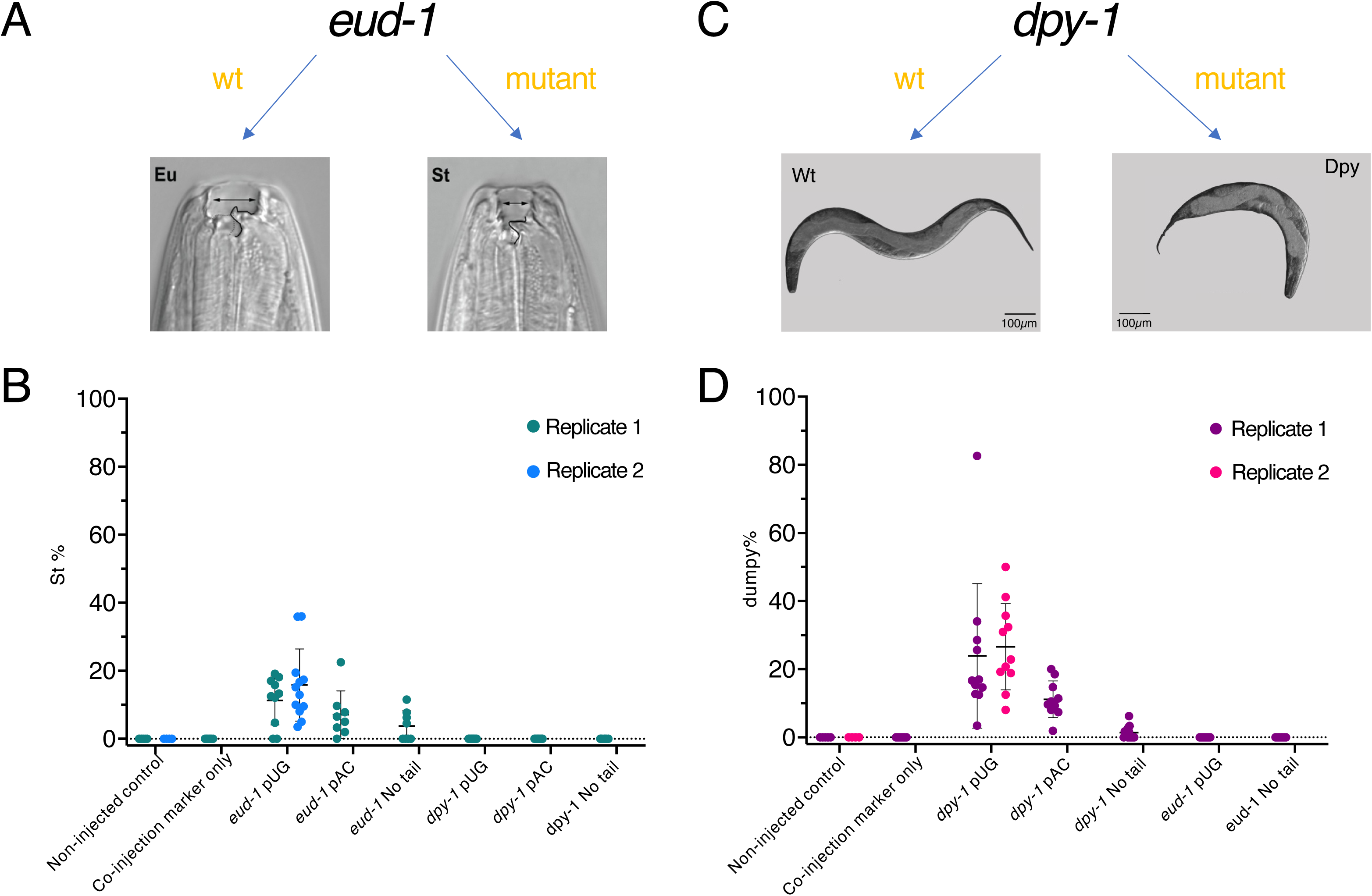
Targeting *eud-1* and *dpy-1*. A and B illustrate the expected phenotypes. A: mouth openings of a eurostomatous (Eu, wide mouth with two teeth) and a stenostomatous (St, narrow mouth with only one tooth) adult *P. pacificus* hermaphrodite. The images are reproduced from (Dardiry et al., 2023) under the CC BY 4.0 license. B: a wild type (Wt) and a dumpy (Dpy, genotype *dpy-1(sy304)*) adult *P. pacificus* hermaphrodite. The indicated RNA was injected into PS312 *P. pacificus* young adult hermaphrodites along with a plasmid encoding RFP under the control of the *eft-3* promoter as a marker for successful injection (co-injection marker). The entire progeny of injected animals that produced any RFP positive progeny was scored. On the Y-axis the % of phenotypically mutant (stenostomatous (St) in C and Dpy in D) progeny of the injected worms is indicated. Every dot represents the brood of one injected animal. The means plus/minus one standard deviation are indicated.

Next, we targeted the RFP transgene in the strain RS3832, which carries a *daf-1::Turbo-rfp* transgene in the PS312 genetic background (Fig. 2). Injection of poly(UG)-tailed RNA led to worms with strongly reduced red fluorescence in the progeny of injected hermaphrodites with moderate to high penetrance. Again, also poly(AC)-tailed and non-tailed RNAs induced RFP silencing, with comparable penetrance to poly(UG)-tailed RNA in the case of poly(AC)-tailed RNA but with clearly lower penetrance in the case of non-tailed RNA. Thus, for three different genes, we have demonstrated that poly(UG)-tailed RNAs can be used to specifically knock down individual genes in *P. pacificus*. Since the penetrance of gene silencing was the highest for the RFP transgene, for the following experiments, we concentrated on knocking down the RFP in RS3832.

**Fig. 2:**
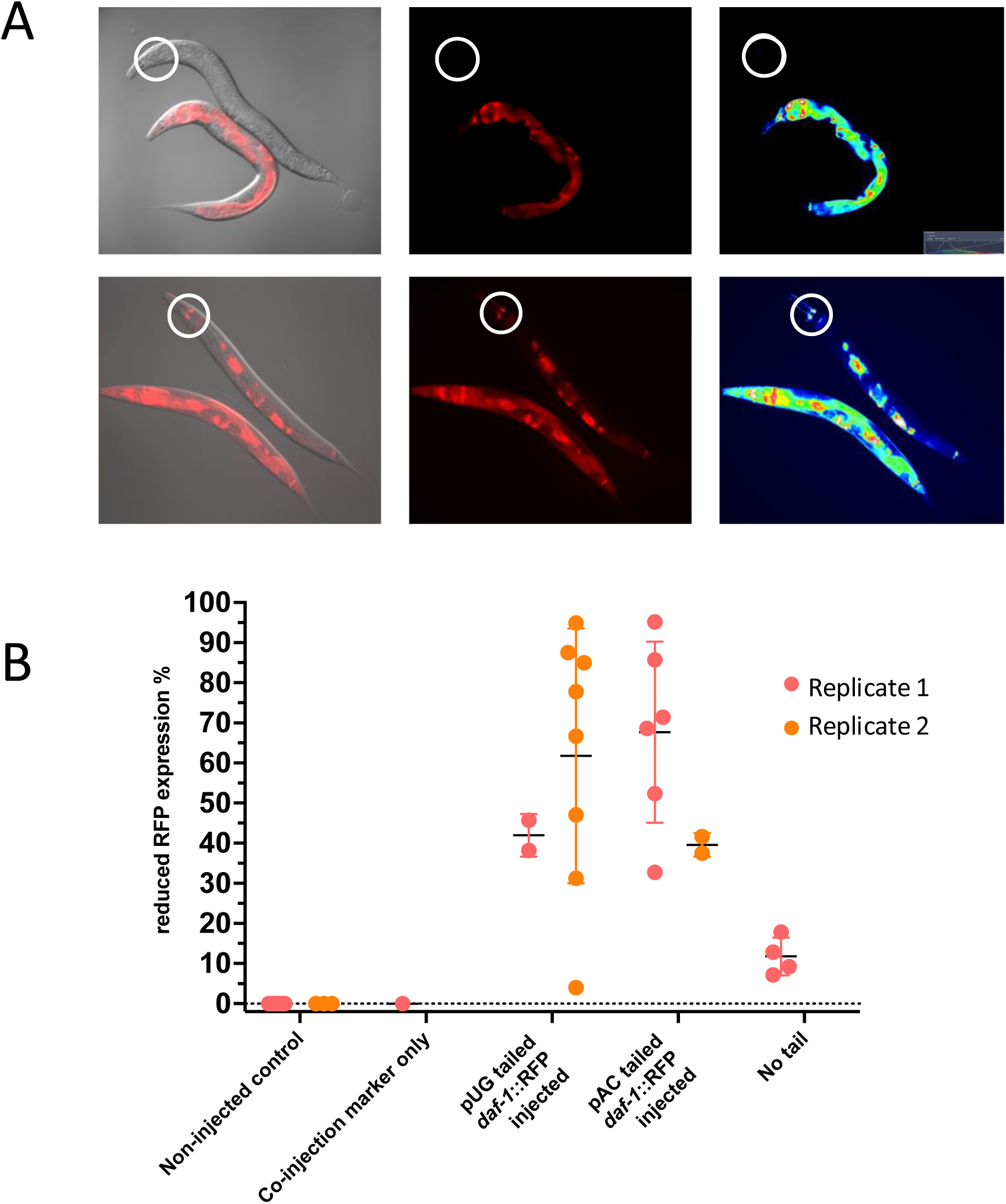
Targeting the multi copy *daf-1::rfp* transgene in RS3832. A: Illustration of the range of the phenotype observed. In the top and the bottom panels, one worm that is the progeny of a poly(UG)-RNA injected mother is shown above a control worm. The central panels show the red channel, the left panels show merged fluorescent and DIC images and the right panels show false colored images of the red channel, indicating signal intensity. The worms scored as “reduced RFP expression” were in the range from the one at the top to the one at the bottom panel. Two cells whose signal never disappeared completely are circled. B: The indicated RNA was injected into RS3832 *P. pacificus* young adult hermaphrodites along with a plasmid encoding GFP under the control of the *eft-3* promoter as a marker for successful injection (co-injection marker). The entire progeny of injected animals that produced any GFP-positive progeny was scored when they were young adults. On the Y-axis, the % of worms with reduced RFP for each brood is given. Each dot represents the brood of one injected animal. The means plus/minus one standard deviation are indicated. Notice that the experiment was reproduced later multiple times with independently prepared poly(UG)-tailed and non-tailed RNAs in the context of the experiments shown in Fig. 4.

### Gene silencing is dependent on *Ppa-rde-1*

In contrast to the findings of (Shukla et al., 2020) in *C. elegans*, in our experiments, injection of poly(AC)-tailed and non-tailed RNAs also induced gene silencing. To exclude that the silencing effects were caused by unavoidable small amounts of double-stranded RNA produced during the synthesis of the injected RNAs, we followed the strategy of (Shukla et al., 2020) to use *rde-1* mutants. In *C. elegans*, these mutant worms are resistant to RNAi by injection of long double stranded RNA due to deficiency in an earlier step of the pathway, but still susceptible to gene silencing by injection of poly(UG) tailed RNA (Shukla et al., 2020).

In *P. pacificus* we identified three *rde-1* orthologs (Fig. 3). One (protein sequence accession number KAF8384843) had been annotated as *Ppa*-RDE-1 in the databases (encoded by gene model PPA37777 on chromosome I, Fig. 3 and http://www.pristionchus.org - Genome Browser) and is the best reciprocal BLASTP hit with *Cel*-RDE-1. We will refer to this gene as *Ppa-rde-1.1*. The two other paralogs, protein sequence accession numbers KAF8358727 and KAF8358512 are encoded by two genes (gene models PPA00739 and PPA36770) located adjacent to each other in head-to-head arrangement on chromosome III. We will refer to these two genes as *Ppa-rde-1.2* (encoding KAF8358727) and *Ppa-rde-1.3* (encoding KAF8358512) (for the full argument of orthology assignment see Materials and Methods and Fig. 3). We generated CRISPR induced mutations in the *rde-1* paralogs in the RS3832 background (Tables 1 and 2, Suppl. Table 1, see Materials and Methods). As described for *C. elegans rde-1* mutations, (Grishok et al., 2005; Kim et al., 2005) we noticed an increase in the expression of the transgene in triple mutants, suggesting that the *rde-1* orthologs are involved in limiting the expression from multi-copy transgenes as in *C. elegans*. Other than this, these mutants did not show any obvious phenotype. All single and double mutants we tested still showed strongly reduced RFP upon poly(UG)-tailed RNA (Fig. 4, panel b-d) and the effect of non-tailed RNA was not abolished (Fig. 4, panel e and f). However, in the triple mutant, the poly(UG)-tailed RNA had no silencing effect (Fig. 4, panel a), suggesting that either, different from *C. elegans*, *rde-1* function is required also for the poly(UG)-tailed RNA dependent RNAi amplification mechanism or that the silencing we observed was due to a different mechanism.

**Fig. 3:**
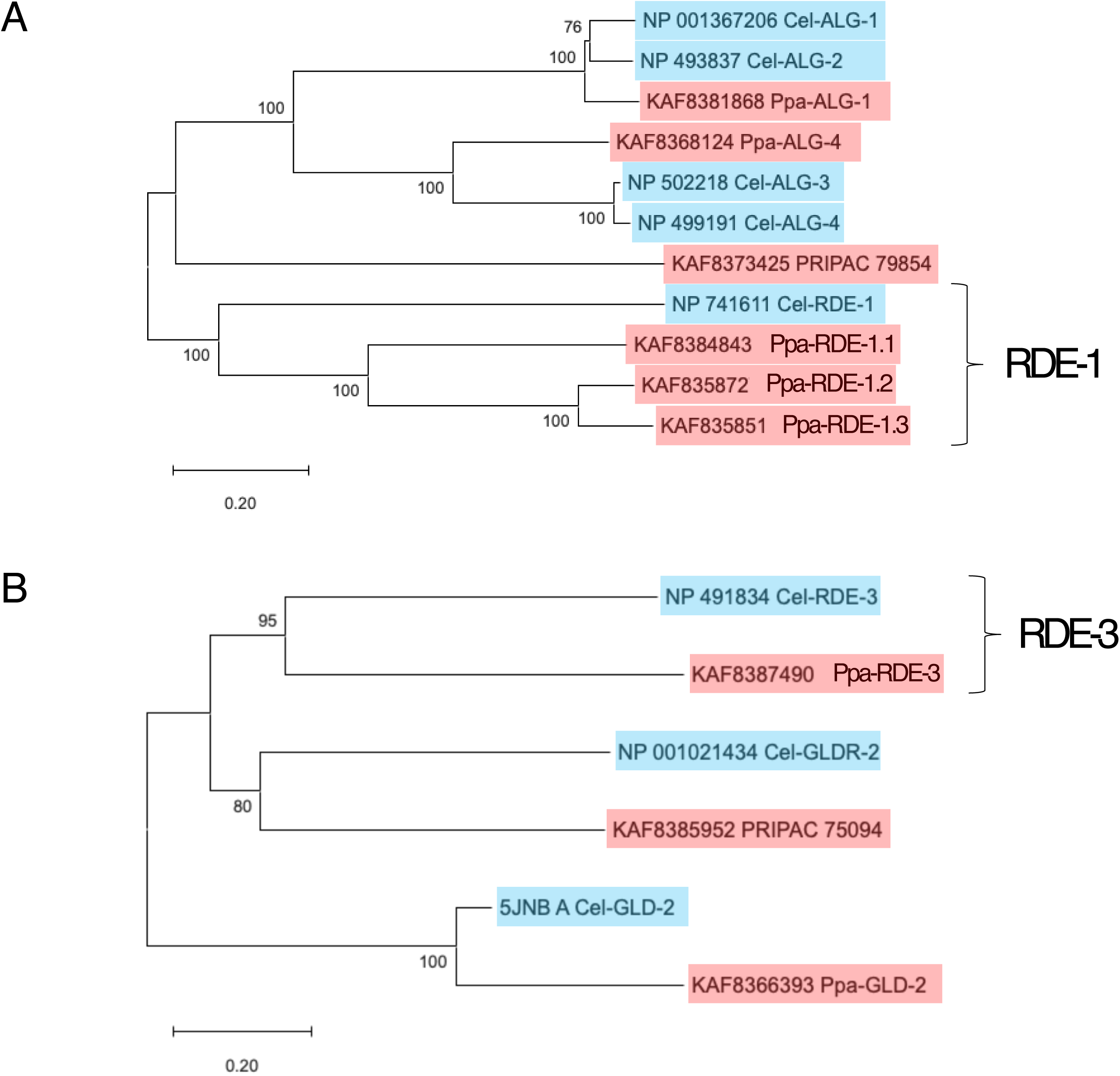
Orthology relationship of *C. elegans* and *P. pacificus* RDE-1. (A) and RDE-3 (B). Neighbor joining trees of *P. pacificus* and *C. elegans* RDE-1 (A) and RDE-3 (B) related genes. Bootstrap values based on 1000 replicates are shown next to the branches. The trees are drawn to scale in units of the number of amino acid substitutions per site. For details on the selection of genes included and for tree reconstruction, see Materials and Methods. Maximum Likelihood trees were also calculated and showed the same topology. GeneBank accession numbers followed by the gene names are given. All gene names other than RDE-1 and RDE-3 are as annotated in WormBase (for *C. elegans*) or Pristionchus.org (for *P. pacificus*) on March 24^th^ 2025. *Ppa-rde-3* was annotated as *Ppa-mut-2* (in *C. elegans*, *mut-2* is a synonym for *rde-3*).

**Fig. 4:**
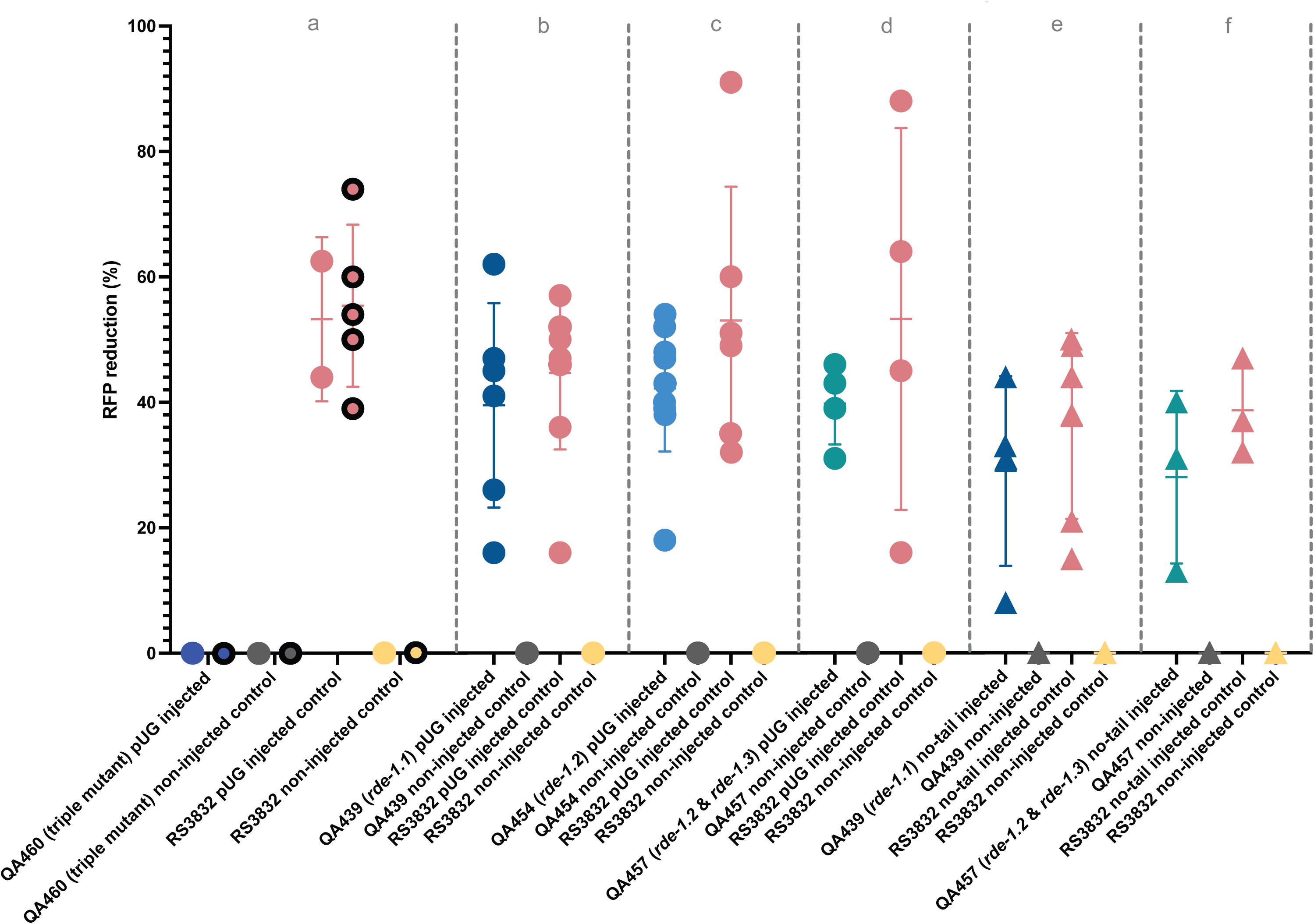
Silencing is dependent on RDE-1. Young hermaphrodites of the indicated strains were injected with poly(UG)-tailed RNA (a-d) or non-tailed RNA (e,f) along with a plasmid encoding GFP under the control of the *eft-3* promoter as a marker for successful injection (co-injection marker). The entire progeny of injected animals that produced any GFP-positive progeny was scored when they were young adults. On the Y-axis, the % of worms with reduced RFP for each brood is given. Each dot represents the brood of one animal in an experiment with poly(UG)-tailed RNA; each triangle represents the brood of one animal in an experiment with non-tailed RNA. The means plus/minus one standard deviation are indicated. Each panel was generated in a different injection session. For a (triple mutant), two injection sessions were done. Control experiments were done in every injection session. The RNAs were synthesized in different/multiple *in vitro* reactions from the ones used in Fig. 2.

**Table 1:**
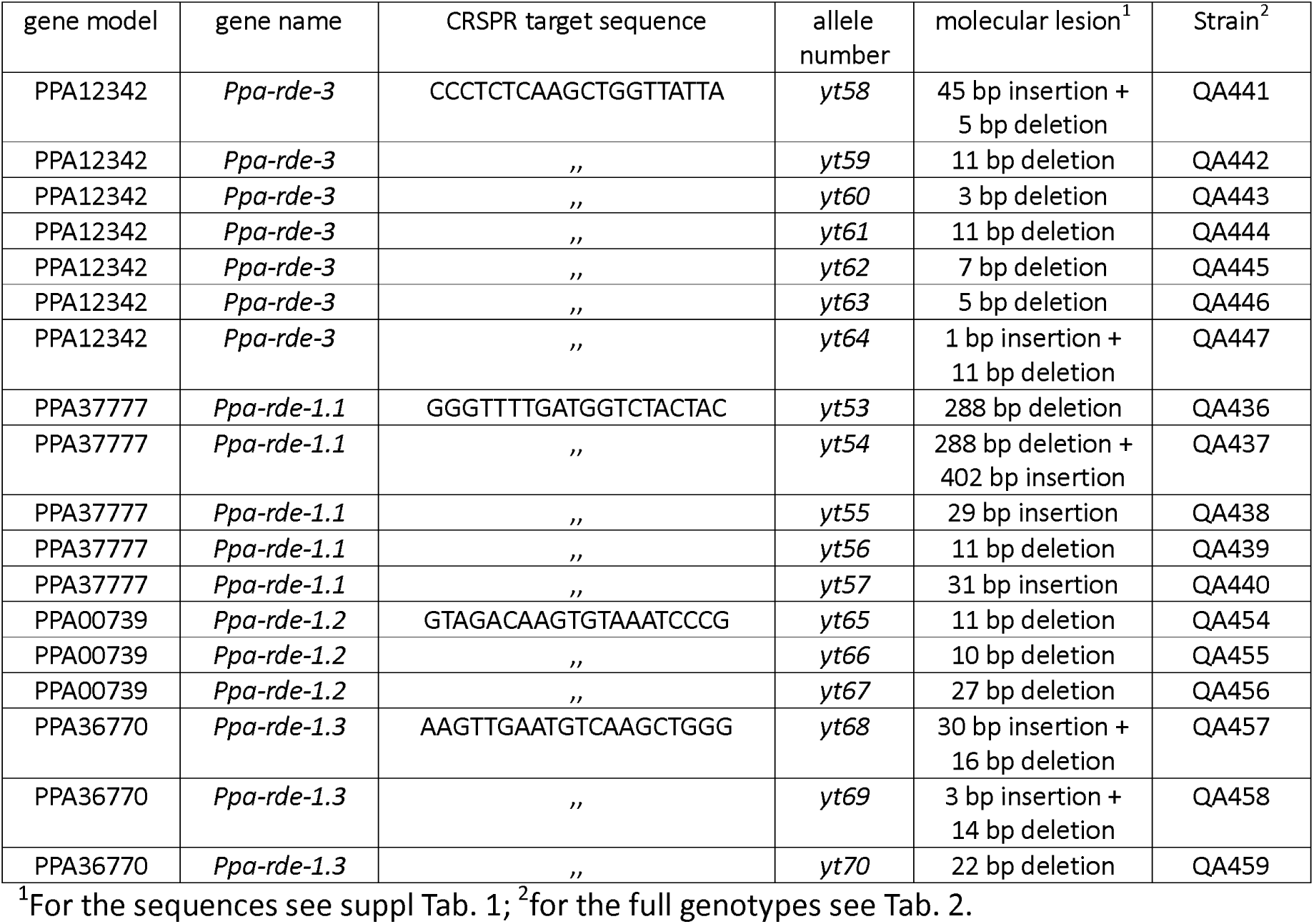
CRISPR/Cas9 induced mutant alleles in Ppa-rde-1.X and Ppa-rde-3.

**Table 2:** Full genotypes of P. pacificus strains used.

| Strain number | full genotype<sup>1</sup> |
| --- | --- |
| PS312 | wt |
| RS3832 | *tuEx333 [daf-1p::TurboRFP]*<sup>1</sup> |
| QA441 | *Ppa-rde-3(yt58)/+; tuEx333 [daf-1p::TurboRFP]* |
| QA442 | *Ppa-rde-3(yt59)/+; tuEx333 [daf-1p::TurboRFP]* |
| QA443 | *Ppa-rde-3(yt60)/+; tuEx333 [daf-1p::TurboRFP]* |
| QA444 | *Ppa-rde-3(yt61)/+; tuEx333 [daf-1p::TurboRFP]* |
| QA445 | *Ppa-rde-3(yt62)/+; tuEx333 [daf-1p::TurboRFP]* |
| QA446 | *Ppa-rde-3(yt63)/+; tuEx333 [daf-1p::TurboRFP]* |
| QA447 | *Ppa-rde-3(yt64)/+; tuEx333 [daf-1p::TurboRFP]* |
| QA436 | *Ppa-rde-1.1(yt53); tuEx333 [daf-1p::TurboRFP]* |
| QA437 | *Ppa-rde-1.1(yt54); tuEx333 [daf-1p::TurboRFP]* |
| QA438 | *Ppa-rde-1.1(yt55); tuEx333 [daf-1p::TurboRFP]* |
| QA439 | *Ppa-rde-1.1(yt56); tuEx333 [daf-1p::TurboRFP]* |
| QA440 | *Ppa-rde-1.1(yt57); tuEx333 [daf-1p::TurboRFP]* |
| QA454 | *Ppa-rde-1.2(yt65); tuEx333 [daf-1p::TurboRFP]* |
| QA455 | *Ppa-rde-1.2(yt66); tuEx333 [daf-1p::TurboRFP]* |
| QA456 | *Ppa-rde-1.2(yt67); tuEx333 [daf-1p::TurboRFP]* |
| QA457 | *Ppa-rde-1.2(yt65) Ppa-rde-1.3(yt68); tuEx333 [daf-1p::TurboRFP]* |
| QA458 | *Ppa-rde-1.2(yt65) Ppa-rde-1.3(yt69); tuEx333 [daf-1p::TurboRFP]* |
| QA459 | *Ppa-rde-1.2(yt65) Ppa-rde-1.3(yt70); tuEx333 [daf-1p::TurboRFP]* |
| QA453 | *Ppa-rde-3(yt61)/+* |
| QA460 | *Ppa-rde-1.1(yt56); Ppa-rde-1.2(yt65) Ppa-rde-1.3(yt68); tuEx333 [daf-1p::TurboRFP]* |

### New tailed RNAs are formed from injected and endogenous RNAs in *P. pacificus*

If the same process as in *C. elegans* is at work, one would expect to see newly formed poly(UG)-tailed RNAs upon injection. We isolated RNA from the injected worms 24 hours after injection and from the progeny of injected worms when they were second and third juvenile stage (notice that *P. pacificus* hatches from the egg shell only as a J2). Following the example of (Shukla et al., 2020), we reverse transcribed the RNA with a primer recognizing the poly(UG)-tail and containing two adaptor sequences. Then, we performed a nested PCR reaction using primers recognizing the 5’ end of the injected RNA and the adaptor sequences (for details see Materials and Methods). In order to avoid that the experiment was dominated by the injected RNA and to increase the chances of identifying poly(UG)-tailed RNAs of different lengths, we separated the PCR products on an agarose gel and isolated and cloned the DNA from different fractions. This rendered this experiment not really quantifiable but merely qualitative. We then sequenced individual clones and found clones with 9 to 26 UG repeats. Notice that the tail length we observed is not indicative for the length of the tail on the RNA molecule the clone was derived from but reflects the place where the reverse transcription primer annealed. For our interpretation we only considered clones with 10 or more UG repeats (Fig. 5A), because the primer used for reverse transcription contained 9 AC repeats and hence the last 9 repeats in the PCR product are derived from the primer. We also manually double checked that there were no DNA encoded UG repeats at the position of the supposed poly(UG) addition site.

**Fig. 5:**
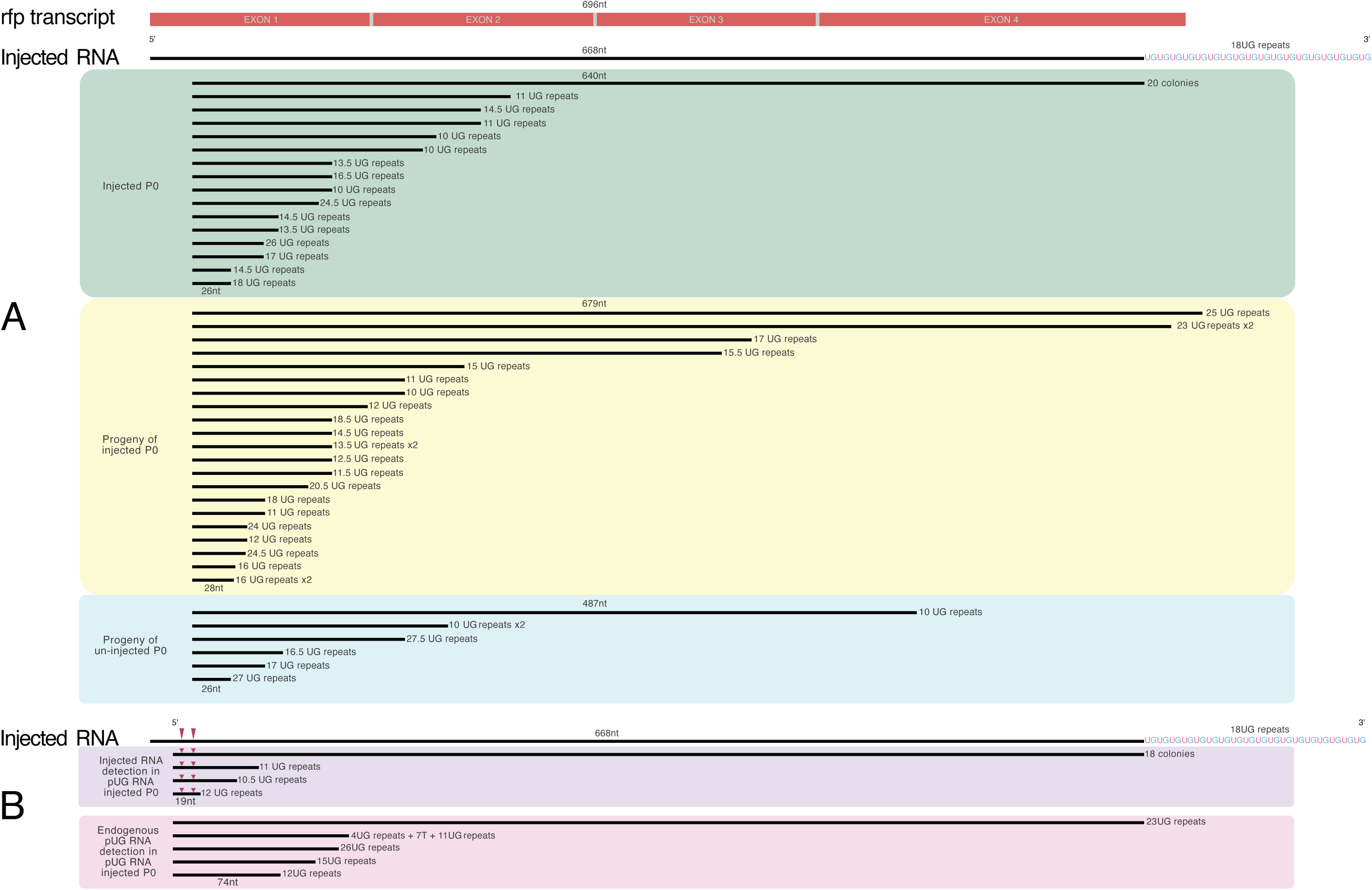
New poly(UG)-tailed RNAs are formed using endogenous and the injected RNA as substrate. A: poly(UG)-tailed RNA was injected into 67 RS3832 *P. pacificus* young adult hermaphrodites. 24h after the injected worms had laid eggs, the worms were collected, pooled and RNA was extracted. The progeny of the injected worms was allowed to hatch and become 2^nd^ and 3^rd^-stage larvae. Of those, all 608 larvae with clearly reduced RFP expression were picked, pooled and used for RNA extraction. An equal number of progenies from un-injected worms were treated in the same way. The RNAs were reverse transcribed with a poly(UG) recognizing primer, amplified according to the “nested PCR” protocol and cloned, as described in Materials and Methods. Clones representing independent reverse transcription events are shown. Only the part of the clone that is derived from the RNA and not from the primers is shown. Notice the two clones from the progeny that extend further 3’ than the injected RNA (see text). B: poly(UG)-tailed RNA marked with two-point mutations was injected into 46 RS3832 *P. pacificus* young adult hermaphrodites. After the injected worms had laid eggs for 24 h, they were pooled and RNA was extracted and analyzed as in A, except that the semi-nested PCR amplification was done using the primers 7363 (biases towards the detection towards the injected RNA) or 7364 (biases towards the detection towards the endogenous RNA) for half of the sample each. The knockdown effect was confirmed in the progeny of these worms. Co-injection marker was included in the injection mix for consistency with the other experiments, but it was not considered in this experiment.

In the sample 24 hours after the injection (Injected P0 in Fig. 5A), as expected, the strongest signal was derived from the injected RNA since 20 out of 35 clones with 10 or more UG repeats contained the full sequence of the injected RNA. The other 15 clones defined ten different poly(UG) addition sites along the RFP sequence. Five of them were represented by more than one clone. These clones had different numbers of UG repeats, indicating that they were independent. In the progeny of the injected worms, we did not detect the injected RNA anymore, but the 21 clones with 10 or more UG repeats defined 14 different poly(UG) addition sites along the RFP sequence of which four were represented by at least two independent clones.

Strikingly, two of these addition sites were upstream of the region covered by the injected RNA (both contained exons 1-3 and a portion of exon 4 of the RFP mRNA, one of them extending into the 3’UTR), indicating that poly(UG) tailed endogenous RFP mRNAs occur naturally. Consistent with the notion that tailed RNAs are naturally produced in *P. pacificus*, we also found six clones derived from the progeny of un-injected P0s with 10 or more UG repeats. These clones defined six different poly(UG) addition sites along the RFP sequence. The presence of poly(UG)-tailed *rfp* RNAs in un-injected worms was also confirmed in adult worms (Fig. 6). In order to determine if the injected RNA was used as a substrate for the production of shorter tailed RNAs, we repeated the experiments with an RNA that, compared with the endogenous one, contained two point mutations at the 5’ end and used primers that either detect preferentially the injected RNA or preferentially the endogenous RNA (see Materials and Methods). We detected tailed products that were shorter than the injected RNA with the point mutations and with the endogenous sequence (Fig. 5B), suggesting, that both, the injected and the endogenous RNA served as substrate for the formation of shorter tailed RNAs.

**Fig. 6:**
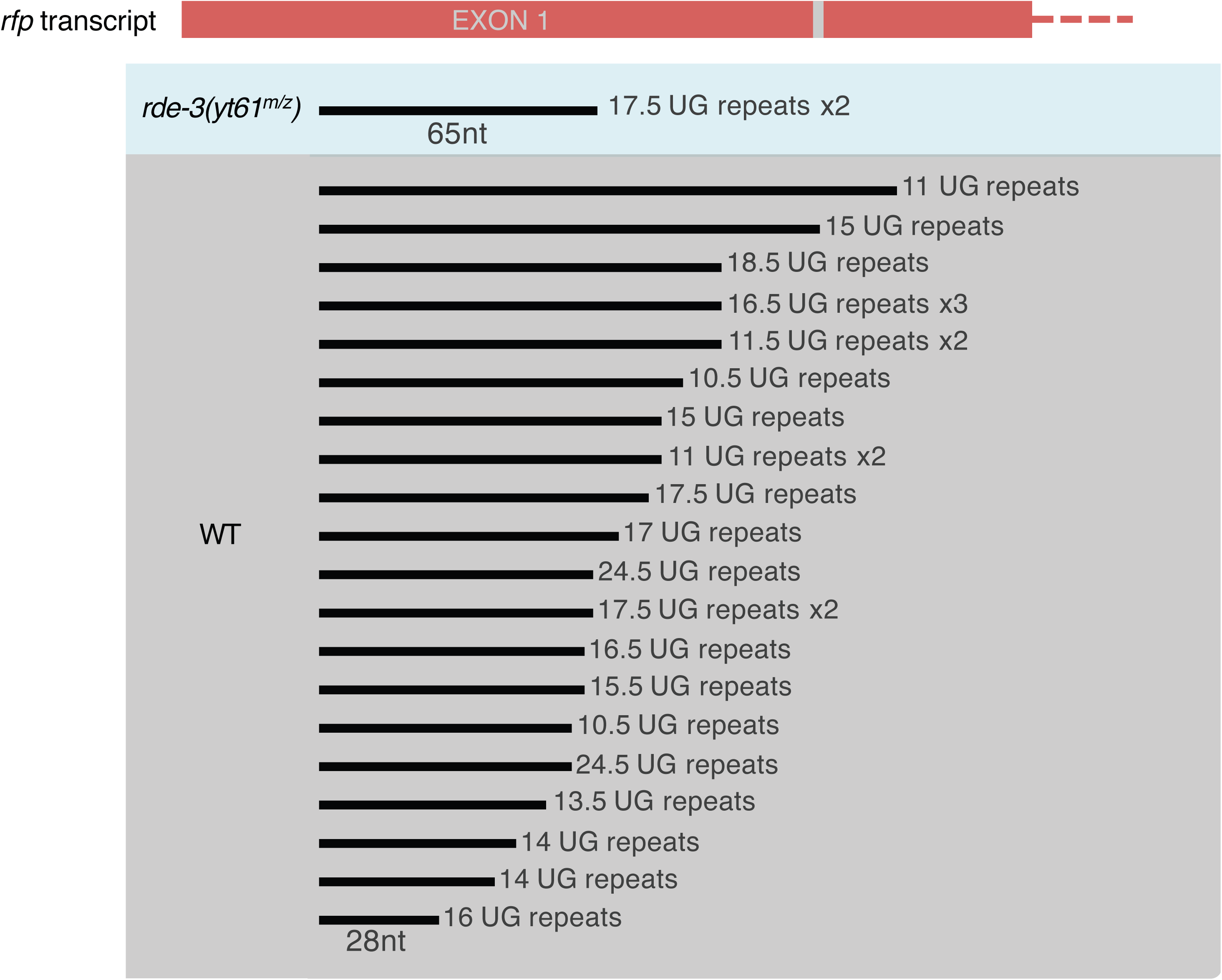
poly(UG) addition is dependent on RDE-3. QA444 segregates *rde-3(yt61)* homozygous, heterozygous and Wt worms. RNA was isolated from 214 adult progenies of *rde-3(yt61)* homozygous mothers and from 214 adult progenies of Wt mothers RNA, and cDNA derived from poly(UG)-tailed RNA was cloned as described in Materials and methods. Clones showing 10 or more UG repeats are shown.

### UG tail addition depends on *Ppa-rde-3*

In *C. elegans*, the polymerase that adds the poly(UG)-tails is encoded by *rde-3*. *P. pacificus* has a one-to-one ortholog of *rde-3* (Fig. 3B, for the full argument for orthology assignment, see Materials and Methods). In order to test if RDE-3 is responsible for the poly(UG) addition in *P. pacificus*, we generated CRISPR induced mutations in this gene in the RS3832 background (Tables 1, 2). All seven *rde-3* alleles isolated are maternal effect sterile (Mes). While most homozygous *rde-3* maternal and zygotic mutant worms are sterile, some mutants lay very few eggs, but the embryos arrest at various stages before hatching (we only ever observed one larva that hatched but did not develop). In order to exclude that the Mes phenotype was aggravated by the stress imposed by the presence of a multi-copy transgene, we backcrossed QA444 (*Ppa-rde-3(yt61)*/+; *tuEx333* [*daf-1p::TurboRFP*]) against PS312 and selected RFP negative worms resulting in QA453. Again, *rde-3(yt61)* showed the Mes phenotype with the very few eggs laid by *rde-3(yt61^m/z^)* being inviable.

We isolated 214 QA444 *rde-3(yt61)* homozygous maternally and zygotically mutant worms (the sterile progeny of maternally rescued homozygous mutants) and tested them for the presence of poly(UG)-tailed *rfp* RNAs (without injection) following the experimental procedure described above. We found only two clones with more than 10 UG repeats (Fig. 6). These two clones were identical (same UG addition site, 17.5 UG repeats) and hence presumably derived from the same original PCR product. In contrast, from an equal number of progeny of QA444 *rde-3(wt)* worms we found 25 clones representing at least 20 different PCR products and 15 different UG addition sites (Fig. 6). Although our assay is not really quantitative, this indicates that RNA tailing is strongly reduced in the absence of RDE-3. The remaining tails may be the product of a small amount of persisting maternal *rde-3* function (protein that remains in the mothers and is passed to the eggs).

### Poly(UG)-tailed RNAs are naturally produced from many endogenous mRNAs

In order to assess whether poly(UG)-tailed RNAs can be naturally generated from mRNAs other than those derived from multi-copy transgenes, we performed RNA 3’- end sequencing. Ribosome and mRNA-depleted RNA from the wt strain PS312 was reverse transcribed with poly(UG) recognizing primers (for details see Materials and Methods). The resulting requesting Illumina data are deposited in the European Nucleotide Archive (Accession number PRJEB9683). We screened for reads that, in addition to a portion that aligned with the *P. pacificus* genome sequence, contained a stretch of (UG) repeats. Since the primer used for the reverse transcription contained 9 UG repeats, we only considered reads that contained 10 or more UG repeats. Out of 8,386,810 total reads that mapped to the genome, 200,087 (2.4%) fulfilled this criterion (Suppl. Tab. 2). In 21,270 (out of a total of 28,896) gene models, we did not find any such reads (73.6%). However, 26.4% of the *P. pacificus* genes produced poly(UG)-tailed RNAs in various amounts. In detail, read number of poly(UG)-tailed RNAs per gene were grouped arbitrarily in very low (1-5 reads; 2,520 genes), low (6-50 reads; 4,356 genes), median (51-100 reads; 466 genes), high (101-500 reads; 266 genes) and very high (501-16,008 reads; 18 genes). At the very top, for instance, were 11 genes with 501 to 1,000 reads, 5 with 1,001 to 2,000 reads and finally one each with 3,331 and with 16,008 reads.

We then filtered out duplicate reads to get an overview of the number of unique reads per gene, which represent independent detection events (reads derived from independent reverse transcription events) (Suppl. Tab. 3). In 5,129 (17.7%) gene models, we found three or more unique reads, of which 1,897 (6.6% of the total) had 10 or more unique reads. 79 models had 100 or more unique reads, among them the top three with 516, 676 and 834. Interestingly, the majority (58.8%) of the 5,129 genes with three or more unique reads fall into one of the five gametogenesis related co-expression modules described by (Athanasouli et al., 2023) and amounted to a considerable fraction of the corresponding module. Specifically, 1,758 genes belong to module 1 (oogenesis 1) and represent 54.9% of the 3,202 genes in the module; 542 genes belong to module 5 (oogenesis 2) and represent 49.3% of the 1,099 genes in the module; 452 genes belong to module 2 (spermatogenesis) and represent 22.7% of the 1,995 genes in the module); 227 genes belong to module 7 (oogenesis 3) and represent 48.1% of the 472 genes in the module; and 36 genes belong to module 23 (oogenesis 4) and represent 50.0% of the 72 genes in the module. We then visually inspected the randomly selected gene models to determine how the reads are distributed over the gene models. Consistent with the hypothesis that a mechanism as described by (Shukla et al., 2020) for *C. elegans* is also at work in *P. pacificus*, the reads aligned to multiple places along the CDS of the gene models.

We wondered if poly(UG)-tailed RNAs exist also in *Strongyloides* spp., a genus of parasitic nematodes in a different clade (clade IV, Blaxter et al. 1998) than *C. elegans* and *P. pacificus* (both in clade V). Since we had found that in *P. pacificus* the injected poly(UG)-tailed RNA served as substrate for the formation of new poly(UG)-tailed RNAs, we injected the same poly(UG)-tailed *rfp* RNA into the gonads of *Strongyloides stercoralis* free-living females and isolated RNA from all injected animals (pooled) 24 h after injection. A co-injection marker (designed for *S. stercoralis*) was included in the injection mix for consistency with the other experiments, but it was not considered in this experiment. In four independent experiments (injecting 44, 29, 28 and 33 animals) with tailed RNAs from two independent in vitro transcription reactions, we detected the injected RNA, but failed to detect RNAs with other poly(UG) addition sites. Next, we performed RNA 3’ end sequencing with RNA extracted from *Strongyloides ratti* and *S. stercoralis*.

In *S. ratti,* we also detected tailed RNAs with 10 or more UG repeats that mapped to multiple, but many fewer genes, compared with *P. pacificus.* We found reads mapping to 7.6% (995) genes of *S. ratti*. As seen in *P. pacificus*, the number of tailed reads per gene also varied. The majority of the tailed RNAs had a very low (1-5 reads; 783 genes), low (6-50 reads; 190 genes), median (51-100 reads; 8 genes) and high (101-1,000 reads; 9 genes) number of reads, but the five top genes had over 1,000 copies of tailed RNA (Suppl. Tab. 4).

In only 212 (1.6%) gene models, we found three or more unique reads, of which 53 had 10 or more unique reads (Suppl. Tab. 5). There were only six gene models with 100 or more reads, with the top three having 169, 639 and 932 reads. Upon visual inspection, we noticed that five of the six gene models with more than 100 unique reads were mitochondrial genes, including the two genes with the highest number of unique reads.

We have also found reads with 10 or more (UG) repeats that mapped to the genome of *S. stercoralis*. However, we only observed tailed RNAs derived from 3.2% (557) of the coding sequences, and for 470 of these the number of reads per gene was a maximum of 5. In only 87 (0.5%) genes, we found more than 5 reads, of which 84 had less than 100 reads. The top six genes produced 53, 67, 82, 100, 273 and 826 copies of the tailed RNA, respectively (Suppl. Tab. 6). In addition, the number of unique reads per gene was low, with 106 genes (0.6%) varying between 3 and 50 unique reads (Suppl. Tab. 7).

The following observations make us believe that most, if not all, of the reads in *Strongyloides* spp. correspond to artefacts. We noticed that the vast majority of reads that aligned with the genome had a large number of mismatches and were therefore not really derived from the sequence they aligned to. Further, if there were multiple unique reads in a particular gene, they tended to map to the same position within the gene. Compared with *P. pacificus*, both, the fraction of reads with 10 or more UG repeats and the number of genes hit by these reads are much lower in *Strongyloides* spp. Our data are not suitable to decide if all these reads are artefacts and poly(UG)-tailed RNAs do not exist in *Strongyloides* spp. or if poly(UG)-tailed RNAs are just less common in this taxon. However, we think that these data can be seen as a maximum estimate of the methodologically introduced background. This supports the notion that most of the poly(UG)-tailed RNAs observed in *P. pacificus* were real.

## Discussion

In *C. elegans*, poly(UG)-tailed RNAs have been shown to be formed in response to the activity of primary siRNAs and piRNAs through the activity of the terminal transferase RDE-3, also known as MUT-2 (Shukla et al., 2020). These RNAs are part of a presumably nematode specific amplification step for the action of siRNAs and piRNAs (Pak and Fire, 2007; Shukla et al., 2020). Injection of poly(UG)-tailed RNAs can be used to experimentally knock down genes (Shukla et al., 2020). Here, we show that a similar mechanism exists in *P. pacificus*. We showed gene knockdown for two endogenous genes and a multi-copy transgene. Different from (Shukla et al., 2020) in *C. elegans*, we observed that injection of non-tailed RNAs or RNAs with a poly(AC)-tail also induced gene silencing. (Shukla et al., 2020) had done their experiments in *rde-1* mutant worms, which are resistant to RNAi initiated by long double-stranded RNAs, which may arise in small quantities in the in *vitro transcription* reaction used to generate the RNAs to be injected. (Shukla et al., 2020) found the silencing effect of poly(UG)-tailed RNAs not to be affected by mutations in *rde-1*. We generated mutations in all three *rde-1* paralogs present in *P. pacificus,* and in our hands, the silencing effect of poly(UG)-tailed RNAs was abolished if all three paralogs were mutant. This may reflect a biological difference between *C. elegans* and *P. pacificus,* but we can also not completely exclude that the silencing we observed is caused, at least in part, by a mechanism different from the one described by (Shukla et al., 2020). However, the fact that we found the injected and the endogenous RNA to be a substrate for the addition of new poly(UG)-tails strongly argues that a similar mechanism is in place in *P. pacificus*. Further, we detected endogenously present poly(UG)-tailed RNAs in thousands of genes (if a cut off of at least three unique reads detected in a gene in our experiment is applied, 5129 genes, which is 17.7% of all predicted genes). Interestingly, more than half (58.8%) of these genes belong to germline-specific co-expression clusters (Athanasouli et al., 2023) and they represent about half of all genes in the four oogenesis clusters and almost a quarter of the genes in the spermatogenesis cluster. There is, however, also a moderate correlation of the number of poly(UG)-tailed RNAs detected and the expression level of genes (germline genes tend to be fairly highly expressed). The notion that poly(UG)-tailed RNAs play an important role in the germline is also supported by the maternal effect sterile phenotype we observed in worms mutant for *Ppa-rde-3*. In *C. elegans*, RDE-3 has been shown to add poly(UG)-tails to RNAs of germline and soma expressed genes.

We failed to find convincing evidence for the existence of a poly(UG)-tailed RNA dependent silencing pathway in *S. stercoralis* or *S. ratti*. However, it should not be firmly concluded that such a pathway does not exist in *Strongyloides* spp. solely based on our negative results. It could be that *Strongyloides* spp. uses a different tail for the same purpose (to elucidate this, a non-tail-biased RNA end seq. could be of help). However, we think that it can be concluded that, if the pathway exists with poly(UG) tails, it is not as widely used in *Strongyloides* spp. as it is in *P. pacificus*. In our RNA end seq experiment that was designed to enrich for poly(UG)-tailed RNAs the proportion of reads with 10 or more UG repeats (and with this more repeats than introduced through the primer used) was about 50 times higher in *P. pacificus* than in the two *Strongyloides* samples and the fraction of genes in which we found three or more unique reads was about 10 times higher in *P. pacificus* compared with *Strongyloides*. While it is not possible to conclude that the pathway is absent from *Strongyloides* spp., the results of the two *Strongyloides* experiments can be taken as maximum estimates for the background detection of poly(UG)-tailed RNAs, which supports that the majority of the reads in the *P. pacificus* experiment were real.

Limitations: Since the RNAi mechanism in *P. pacificus* is poorly characterized we have no mutants of which it is known that they are insensitive to silencing through the primary RNAi pathway that is triggered by long double-stranded RNAs. Although RNAi by the injection of such RNAs is believed to work only in exceptional cases, we cannot exclude that some of the effects we saw were caused by small quantities of double-stranded RNAs that arose in the *in vitro* transcription reaction.

We would like to point out very clearly that our PCR and cloning-based experiments used to detect poly(UG)-tailed RNAs are purely qualitative (demonstrate the existence and identify addition sites) but not quantitative. Although in principle more quantitative, also in the RNA end sequencing experiment, the quantitative information is limited, due to the extensive pre-treatment of the RNA (rRNA and poly(A) RNA depletion) and the PCR amplification step during the library construction.

## Conclusions

Here, we show that in *P. pacificus,* a gene regulatory mechanism that involves poly(UG)-tailed RNAs whose formation is dependent on *Ppa-rde-3* is in place. It is of particular importance in the germ line. Poly(UG)-tailed RNAs can be used to experimentally downregulate genes in *P. pacificus*. Different from *C. elegans*, the knockdown requires the redundant activity of the three *rde-1* homologs. In *P. pacificus*, knockdown through injection of poly(UG)-tailed RNAs works more reliably than RNAi based on the injection of double-stranded RNA, but not as efficiently as RNAi works in *C. elegans*. If the clade IV nematodes *Strongyloides* spp. have a similar mechanism, it is much less widely used than in the clade V nematodes *P. pacificus* and *C. elegans*.

## Materials and Methods

### Worm Strains and Cultures

The genotypes of all *P. pacificus* strains constructed and/or used are given in Table 2. RS3832 is not yet published and was a gift from Wensui Lo (at the time at our institute, now at Northwest A&F University, Yangling, China). The strain carries a multi-copy *daf-1::Turbo-rfp* transgene in the PS312 genetic background and was constructed by Wen-Sui Lo when she was in Ralf Sommer’s lab at our institute. Requests for this strain should be addressed to Wen-Sui Lo or Ralf Sommer. Although the transgene was originally established as an extrachromosomal array, it is transmitted to 100% of the progeny and, in crosses, behaves like a Mendelian locus. Hence, we suspect that the array did integrate into a chromosome. *P. pacificus* was maintained on NGM agar plates at 20°C as described for *C. elegans* by (Stiernagle, 2006).

*S. stercoralis* PV001 (Hunt et al., 2016) was a gift from James Lok (University of Pennsylvania) and had been maintained in Mongolian Gerbils (*Meriones unguiculatus*) in the laboratory since April 2021 as described by (Lok, 2007) and (Nolan et al., 1993) with occasional reversion to frozen stock. *S. ratti* ED321 was a gift from Mark Viney (at the time University of Bristol, now University of Liverpool) and had been maintained in the lab since 2010 in female Wistar rats as described by (Viney et al., 1992) with occasional reversion to frozen stock. All relevant animal welfare regulations (the German “Tierschutzgesetz” and the **"**Verordnung zum Schutz von zu Versuchszwecken oder zu anderen wissenschaftlichen Zwecken verwendeten Tieren) were followed. Permits with respect to animal experimentation (EB 04/20 A and EB 02/23 V), infection protection (AZ: 25-28/5420.11-3.9 / Streit, Adrian) and animal health (AZ: 3STV/9114.51 / MPI Streit) were granted by the “Regierungspräsidium Tübingen". Animal and laboratory facilities are subject to regular inspection by the authorities.

### PCR and sequencing

All PCR primers were purchased from Eurofins Genomics unless otherwise specified. All PCR reactions were done using the Qiagen Taq master mix (201445) or DreamTaq PCR Master Mix (K1081). All primer sequences are listed in Table 3. All PCR products to be sequenced were submitted to Genewiz from Azenta Life Sciences. 0.5-2µl of the PCR product mixed with 1µl of the sequencing primer (stock concentration 10 mM) were sent in a 10µl.

### Construction of the template DNA for *in vitro* transcription

Total RNA (1-2µg) extracted from mixed stage RS3832 using TRIzol (life technologies, 15596026) and following the manufacturer’s instructions was used to synthesize cDNA using superscript II (Invitrogen, 18064-014) with oligo(dT) - (QT796) primers in a 20µl reaction. RNA was heated with 1µl of 10mM dNTPs, 1µl of reverse transcription (RT) primer (oligo(dT) - QT796) and H_2_O to 65 °C for 5 min and immediately chilled on ice. Then the 4µl of 5x first strand buffer, 2µl of 0.1M DTT and 1µl of H_2_O were added to the reaction and incubated at 42°C for 2mins.

Finally, 1µl of Superscript II (or H_2_O for the negative control) was added and incubated for 2hrs at 42°C followed by 15min at 70°C. 1µl of the cDNA was used as template to amplify the target portion of the gene with gene-specific primers in a 10µl reaction:eud-1 (F-7291 & R-7294), dpy-1 (F-7275 & R-7276), rfp (F-7307 & R-7311). The PCR products were sequenced from both sides using the amplification primers. The PCR products were diluted 1:100 and 0.5µl was used as template for a second PCR with primers containing overhang sequences to add the T7 promoter in the 5’ end and the desired tail on the 3’ end (see primer list: 7297-eud-1 F_T7, 7283-dpy-1 F_T7, 7312-rfp F_T7, 7299-eud-1 R_pUG, 7300-eud-1 R_pCA, 7284-dpy-1 R_UG, 7285-dpy-1 R_CA, 7318-rfp R_pUG, 7317-rfp R_pAC and for the no tail PCR the corresponding T7 primer and the non-overhang R primer was used). These products were gel purified on 1% agarose, 1x TAE gels and the DNA was extracted from the gel using the QIAquick Gel Extraction Kit (28706) following the manufacturer’s instructions. The purified PCR products were sequenced from both sides using the amplification primers. As an alternative template for the generation of *rfp* targeting RNAs and to avoid problems with a non-productive alternative splice form of the *rfp*-RNA we found to be present in the worms, the 1^st^ PCR of the *rfp* fragment (without overhangs) was cloned using the TOPO TA cloning kit (Invitrogen 450071 & 450641) and one shot competent cells (Invitrogen C404010) resulting in plasmid pVdR1. The correct sequence was confirmed by sequencing from both sides using the M13 primers.

### Generation of SNP marked RNAs

In order to obtain *rfp* RNA marked with two point mutations, instead of F-7307 we used a forward primer (7344) carrying the SNP and the T7 promoter to amplify the template for *in vitro* transcription. In these RNAs the SNPs are close to the 5’ end such that even very short molecules could be detected. However, for RNA detection, only one primer site was available at the 5’ end, such that the “semi-nested” protocol was used for detection (see below). In order to also have the opportunity for detection with fully nested PCR, we also generated a construct with SNPs a bit further downstream. First, we used primer F-7375, carrying the SNPs for a PCR with R-7311 for 10 cycles. From this PCR reaction, 0.5 µl was used for 20 cycles of PCR with the primer F-7376 (whose 3’ part overlaps with the 5’ part of 7375) and R-7311. This PCR product was cloned as before, resulting in plasmid pVdR2. Then this plasmid was used as the template for the PCR with the overhang primers that introduce the T7 promoter and the tails as described above.

### In vitro transcription of test RNAs

*In vitro* transcription of the constructs described above was performed using the Invitrogen, MEGAscript™ T7 Transcription Kit (AM1333) according to the manufacturer’s instructions with the following modifications. The transcription reaction volume was scaled up to 40µl and the input DNA was increased to 1.5µg. The incubation step was increased to 37°C overnight for a better yield. Turbo DNase treatment at 37°C was done for 15-40 min.

Following the Turbo DNase treatment, the reaction was extracted once with Phenol: Chloroform:Isoamyl Alcohol 25: 24: 1 (PanReac, AppliChem, A0889,0100) and the RNA was precipitated with one volume of isopropanol (ROTH, 7343.1).

The RNA pellet was washed twice with ice-cold 80% Ethanol, air dried and resuspended in 15µl of H_2_O. The RNA concentration was determined by nanodrop analysis of 1 µl of a 1:10 dilution, and 200 ng of the RNA were analyzed on a 10% polyacrylamide, 1x TBE, 8M urea gel.

### Injection procedures

The injection mix consisted of the RNA to be tested and a DNA plasmid encoding a fluorescent protein as a co-injection marker in TE. In the pilot experiment (Suppl. Fig. 1), RNA concentrations of 0.17pmol/µl, 0.5pmol/µl, 1.5pmol/µl and 4.5pmol/µl were tested for *eud-1*. All later experiments were done using 4.5pmol/µl.

For *P. pacificus,* the co-injection markers were plasmids encoding RFP (for experiments targeting *eud-1* or *dpy-1*) or GFP (for experiments targeting *rfp*) under the control of a *Ppa-eft-3* promoter. Requests for detailed maps of these plasmids or the plasmids themselves should be addressed to Hanh Witte or Ralf Sommer. The co-injection markers were injected at 30ng/µl. For *S. stercoralis,* the co-injection marker was 50ng/µl of the commercially available plasmid pAJ50 (Addgen), which encodes RFPmars under the *Sst-act-2* promoter.

The injection mix was micro-injected into the gonads of the young adult worms (P0s). All the P0s were singled out individually on NGM plates. In the case of *S. stercoralis*, 3-5 males were added to each female. The injected P0s were allowed to lay eggs for 24 hrs. Then the adult worms were removed from the plates and the F1 progeny were allowed to develop into the developmental stage desired for the respective experiment.

### Detection of tailed *rfp* RNAs

The worms specified for the particular experiment in the result section were flash frozen in as little H_2_O as possible. Total RNA was extracted using TRIzol (life technologies, 15596026), DNase I treated at RT for 25mins and cleaned and concentrated using Zymogen Clean & Concentrator (R1013) according to the manufacturer’s instructions.

The RNA was reverse transcribed using a tail recognizing primer 7755 with 9UG repeats and 2 adaptors (Ad1 and Ad2) as described above.

For the following PCR reaction, one of three forward primers was used: primer 7307 (for the nested protocol, this primer recognizes the injected and the endogenous RNA equally well), 7363 (for the semi-nested protocol, this primer recognizes the 5’end of the injected RNA including the 10 first nucleotides that are derived from the transcription template and are not present in the endogenous RNA and therefore biases towards the detection towards the injected RNA) or 7364 (for the semi-nested protocol, this primer extends 10 nucleotides further upstream than the injected RNA and therefore biases towards the detection towards the endogenous RNA). A first PCR was done using the desired forward primer and the Ad1 7757 reverse primer. The product was diluted 1:1000, and 1µl was used as template in the second PCR with the Ad2 reverse primer 7758 and either the nested (7319, nested protocol) or the same forward primer as in the first round of PCR (semi-nested protocol). The resulting PCR products were separated on a 1% agarose, 1x TAE gel. The size range between 1000 bp and 100bp of the lane was divided into multiple pieces. The DNA was extracted using the QIAquick Gel Extraction Kit (cat no. 28706) according to the manufacturer’s instructions and cloned using the Invitrogen TOPO TA cloning kit (450071 & 450641). All (if there were few) or 20 colonies per fraction were tested by colony PCR using the standard M13 primers and analyzed on a 1% agarose, 1x TAE gel. Clones that appeared to be derived from primer dimers were discarded. The remaining clones were sequenced using the standard M13 primers.

### 4.7. Sequence analysis

Sanger sequences were manually checked using the SnapGene software (SnapGene® software (from Dotmatics; available at snapgene.com) in order to determine the position to which the poly(UG)-tail was added. Only clones with 10 or more TG repeats were considered (9 repeats are in the primer used for reverse transcription). If the position of the UG tail was a TG rich site, the sequence was disregarded. For Illumina sequences, adapter sequences were removed by the cutadapt tool (version 4.4 with options -a TGGAATTCTCGGGTGC -A GATCGTCGGACTGTAG -m 20:20). Trimmed read pairs with minimum length of 20 nucleotides were then aligned against the *P. pacificus* genome (version El Paco) by the STAR alignment tool (version 1.7.10b with option --alignIntronMax 10000) (Dobin et al. 2013; Rödelsperger et al. 2017). Duplicate reads were removed by the samtools rmdup program and the binary alignment file (.bam) was converted into text format (.sam) using the samtools view command (version 0.1.18) (Li et al. 2009). Based on the cigar string and the nucleotide sequence in the .sam file, we extracted reads containing a portion that aligned to the genome and also contained 10 or more unaligned TG repeats. Quantification of UG-containing reads against the *P. pacificus* gene annotations (version El Paco gene annotation version 3) was performed using the featureCounts program as implemented in the R subreads package (version 4.0.0) (Liao, Smyth, and Shi 2014; Athanasouli et al. 2020). For further manual inspection, the extracted reads were converted into .bam format using the samtools view command and then visualized in the genome browser.

### Microscopy

Phenotyping for all 3 genes was done when the worms were in the young adult stage. The morphological phenotypes (mouth form and Dpy) were scored while the worms were on NGM plates, using Zeiss Discovery V20 and Olympus SZX10 high-power dissecting scopes with illumination from underneath. RFP and GFP fluorescence were assessed while the worms were on NGM plates, using a Leica M205 FCA fluorescence dissecting scope. For higher magnification images, a Zeiss Imager Z1 fitted with an Axiocam 506 mono camera was used.

### Identification of *P. pacificus* RDE-1 homologs

All BLAST analyses were performed at NCBI (https://blast.ncbi.nlm.nih.gov/Blast.cgi) and repeated on March 24^th^ 2025. The protein sequence of *C. elegans* RDE-1 was retrieved from the databases (accession number NP_741611.1). BLASTP analysis against *C. elegans* and *P. pacificus* protein sequences revealed NP_741611.1 and KAF8384843.1 as best reciprocal BLASTP hits. KAF8384843.1 had been annotated as *P. pacificus* RDE-1 in the database entry. Both, NP_741611.1 and KAF8384843.1, when used as bait for BLASTP searches against *P. pacificus*, returned the same six top hits with a large increase of the e-values between the third and the fourth hit (accession numbers KAF8384843.1 (annotated as *Ppa*-RDE-1), AF8358727.1 (PRIPAC_93722), KAF8358512.1 (PRIPAC_93507), KAF8381868.1 (annotated as *Ppa*-ALG-1), KAF8373425.1 (PRIPAC_79854), KAF8368124.1 (annotated as *Ppa*-ALG-4).

When used as bait for BLASTP searches against *C. elegans*, they returned the same five top hits with a large increase in the e-values between the first and the second hit (accession numbers NP_741611.1 (*Cel*-RDE-1), NP_001367206.1 (*Cel*-ALG-1), NP_493837.1 (*Cel*-ALG-2), NP_502218.1 (*Cel*-ALG-3), NP_499191.1 (*Cel*-ALG-4). From these 11 sequences, a Neighbour Joining (NJ) tree (Fig. 3) was reconstructed as described below. A Maximum likelihood tree was also computed and showed the same topology as the NJ tree.

### Identification of *P. pacificus* RDE-3

All BLAST analyses were performed at NCBI (https://blast.ncbi.nlm.nih.gov/Blast.cgi) and repeated on March 24^th^ 2025. The protein sequence of *C. elegans* RDE-3 was retrieved from the databases (accession number NP_491834.1). BLASTP analysis against *C. elegans* and *P. pacificus* protein sequences revealed NP_491834.1 and KAF8387490.1 as best reciprocal BLASTP hits. KAF8387490.1 had been annotated as *P. pacificus* MUT-2 (a.k.a. RDE-3, see introduction) in the database entry. Both NP_491834.1 and KAF8387490.1, when used as bait for BLASTP searches against *P. pacificus*, returned the same three top hits with a large increase in the e-values between the first and the second hit (accession numbers KAF8387490.1 (annotated as *Ppa*-MUT-2), KAF8385952.1 (PRIPAC_75094), KAF8366393.1 (annotated as *Ppa*-GLD-2). When used as bait for BLAST searches against *C. elegans*, they returned the same three top hits with a large increase in the e-values between the first and the second hit (accession numbers NP_491834.1 (*Cel*-RDE-3), NP_001021434.1 (*Cel*-GLDR-2), 5JNB_A (*Cel*-GLD-2)). From these 6 sequences, a Neighbour Joining (NJ) tree (Fig. 4) was reconstructed as described below. A Maximum likelihood tree was also computed and showed the same topology as the NJ tree.

### Phylogenetic analysis and tree reconstruction

The sequences were aligned with Muscle in MEGA12 (Kumar et al., 2024; Stecher et al., 2020) using default settings. The following information was retrieved from the captions to the trees provided by MEGA. The evolutionary history was inferred using the Neighbor-Joining method (Saitou and Nei, 1987). The optimal trees with the sum of branch length = 4.170 (tree in Fig. 3A) and 3.216 (tree in Fig. 3B) are shown in Fig. 3. The percentage of replicate trees in which the associated taxa clustered together in the bootstrap test (1,000 replicates) are shown next to the branches (Felsenstein, 1985). The trees are drawn to scale, with branch lengths in the same units as those of the evolutionary distances used to infer the phylogenetic tree. The evolutionary distances were computed using the Poisson correction method (Zuckerkandl and Pauling, 1965) and are in the units of the number of amino acid substitutions per site. The pairwise deletion option was applied to all ambiguous positions for each sequence pair resulting in a final data set comprising 1454 positions for the tree in Fig. 3A and 995 positions for the tree in Fig. 3B. Evolutionary analyses were conducted in MEGA12 (Kumar et al., 2024; Stecher et al., 2020) utilizing up to 7 parallel computing threads.

### CRISPR knockout mutants of *rde-1* and *rde-3*

Knockout mutants were generated via CRISPR/Cas9 mutagenesis following (Witte et al., 2015), with modifications described in “<u>Co-Crispr injection protocol</u> by Hanh Witte (version 2019-10-23)” in the “protocols” section of Pristioncus.org (http://www.pristionchus.org/download/protocol_cocrispr_2019_10.pdf). The co-injection marker used was the same *Ppa-eft-3::rfp* construct as used for the RNA injection experiments. The sequences targeted are listed in Table 1.

### Detection of poly(UG)-tailed RNA detection in *P. pacificus*, *S. ratti* and *S. stercoralis* via [poly(UG) biased] RNA sequencing

Total RNA was extracted from mixed-stage cultures of *P. pacificus* (PS312) and free-living *S. ratti* (ED321) and *S. stercoralis* (PV001) animals as described above. The RNA samples were depleted from ribosomal RNA and poly(A) RNA using the *P. pacificus* (Ribo-Seq) riboPOOL (siTOOLs BIOTECH, dp-K006-000082) kit (for *P. pacificus*), the *S. ratti* (Ribo-Seq) riboPOOL (siTOOLs BIOTECH, dp-K006-000106) (for *S. ratti* and *S. stercoralis*), and the Poly A riboPOOL (siTOOLs BIOTECH, dp-K012-34) (all three samples) according to the manufacturer-provided protocols as follows. The rRNA depletion RP and poly A RNA depletion reagents were mixed 1:10 and 1µl of this mix was used in the hybridization step with total RNA (14µl of 2µg) and hybridizing buffer (RNase inhibitor was skipped). The mixture was incubated at 68°C for 10mins and allowed to cool down slowly to 37°C. 80µl of prepared beads were mixed with the hybridized riboPOOL and total RNA and incubated at 37°C for 15mins, followed by an 50°C incubation for 5 minutes. The tube with the mix was placed on a magnetic rack for two minutes to pellet the beads and the supernatant with the rRNA and poly A-depleted RNA was kept. The last step was repeated with the supernatant to get rid of remaining tract amounts of beads. The RNA was purified using the clean-up beads purification provided with the kit as per the manufacturer’s instructions.

The rRNA and poly A depleted RNAs were analyzed on an Agilent Bioanalyser 2100 along with an aliquot of the total RNA to assess the depletion quality. The *S. stercoralis* RNA sample contained some remaining ribosomal RNA compared with the other two. This might be because no rRNA species-specific kit is currently available for *S. stercoralis* and the one designed for *S. ratti* was not fully effective. The RNAs were precipitated overnight using x3 RNA volumes of ice cold 100% EtOH, 10% of RNA volume of 3M NaOAc and 1µl of Glycol Blue. The pellets were washed twice with 70% EtOH, air dried and resuspended in 5µl of nuclease-free water.

To remove the 5’ cap and generate 5’ mono-phosphate RNAs, the RNAs were treated with CapClip Pyrophosphatase (CCP, Biozym) as follows: 2µl of x10 CCP buffer, 2.5µl (1 U/µl) of CCP enzyme (freshly diluted in water from the 5 U/µl stock), 10.5µl water and the 5µl of RNA from the previous step were mixed and incubated for 1hr at 37°C followed by a cleaning and concentration step using the Zymo Research RNA clean and concentrator-5 kit (R1013). The RNA was eluted in 6.5µl of water.

Ligation of the 5’ adaptor was achieved by incubating 1µl of 5µM RA5 adaptor (see the primer list) with the RNA at 70°C for 2min and then transferred immediately to ice. Then 0.8µl of x10 RNA ligase buffer (NEB, B0216S), 0.8µl of T4 RNA ligase 1 (NEB, M0204S) and 0.8µl of 10mM ATP were added and the mix was incubated at 37°C for 2hrs.

The 5’ adaptor ligated RNA was reverse transcribed using a primer recognizing the poly(UG)-tail (7485): 1µl of the 4µM 7485 primer was mixed with 8µl of the RNA, incubated at 70°C for 2mins and immediately put on ice. Then 4µl of 5x 1^st^ strand buffer, 4µl of 10mM dNTP, 1µl of 0.1M DTT, 1µl of superscript III (Invitrogen, 18080-044) and 1µl of water were added and incubated at 50°C for 1hr followed by heat inactivation at 70°C for 15mins.

Then the libraries were prepared using the Illumina TrueSeq kit (15016912). Based on a small-scale trial PCR, we decided to use 21, 25 and 24 amplification cycles for *S. stercoralis*, *S ratti* and *P. pacificus,* respectively. For each sample, three 50µl reactions were set up in parallel. The 3 reactions were pooled, 2µl of the *Strongyloides* and 3µl of the *P. pacificus* were run in a 1% agarose gel to check the quality. The pooled reactions were precipitated overnight using three volumes of 100% ice-cold EtOH. After the centrifugation, the pellets were washed twice with 70% EtOH, air dried and was resuspended in 25µl of water and 1µl was diluted 1:10 and analyzed on Qubit (Thermo Fisher) and an Agilent Bioanalyser 2100.

The libraries were fractionated/size selected using the BluePippin system (1.5% gels with internal marker) to obtain 2 libraries of the same sample with two size ranges: 300-600bp and 600-1500bp. 2.5nM of the cDNA library from each of the 3 species were pooled to result in 2 pools of the 2 library sizes. The libraries were submitted for Illumina sequencing on an Illumina Nexseq 2000 instrument.

## Supporting information

Primer list

## Acknowledgements

We thank Dafne Ibarra and Natali Betz for methodological support and Ralf J. Sommer and Wen-Sui Lo for helpful discussions and *Pristionchus* strains.

---

**Suppl. Fig. 1 Pilot experiment:** poly(UG) tailed RNA targeting *eud-1* at the indicated concentrations was injected into PS312 *P. pacificus* young adult hermaphrodites along with a plasmid encoding RFP under the control of the *eft-3* promoter as a marker for successful injection. The entire progeny of injected animals that produced any RFP positive progeny was scored. Since the injections had to be done sequentially, at every time worms were picked for injection, control worms were also picked from the same culture. Each dot represents the progeny of one mother. %ST: % stenostomatous individuals.

