## Supplementary figures and images for "Poly(UG)-tailed RNAs are involved in the control of thousands of genes predominantly in the germline in *Pristionchus pacificus*"

### Supplementary Figure 1

Suppl. Figure 1

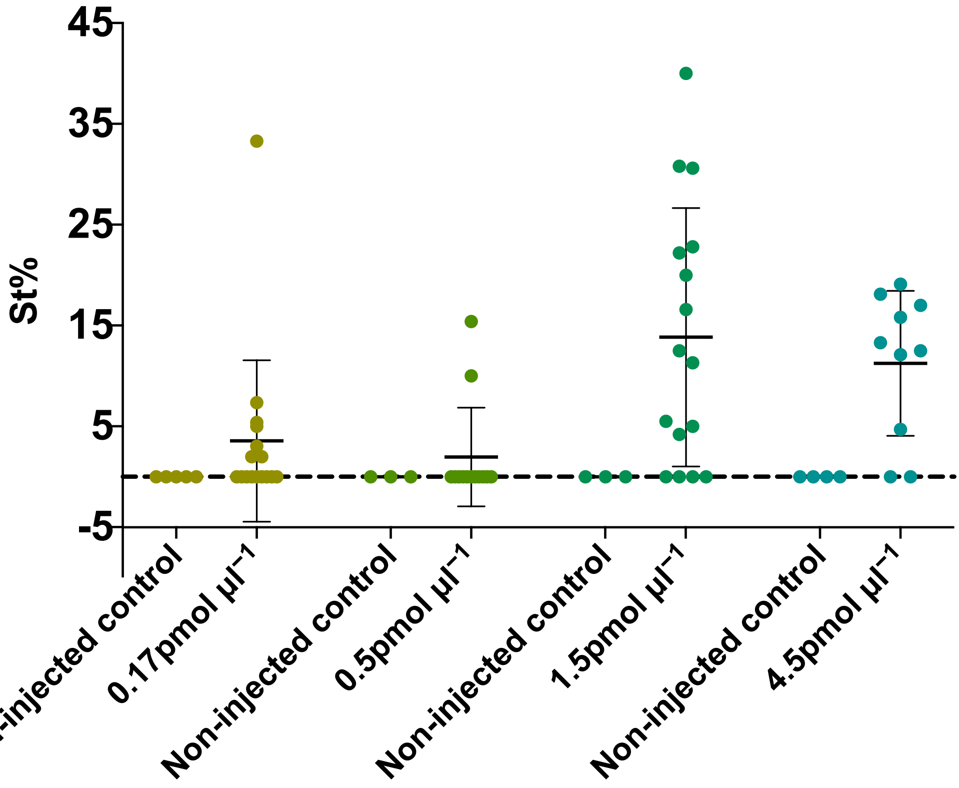
