## Supplementary Table 1 for "Poly(UG)-tailed RNAs are involved in the control of thousands of genes predominantly in the germline in *Pristionchus pacificus*"

Suppl_Table 1: CRISPR/Cas9 induced mutant alleles in *Ppa-rde-1.X* and *Ppa-rde-3*

| Gene | allele | molecular lesion | Sequence (insertions in blue and deletions in red) |
| --- | --- | --- | --- |
| *rde-3* | *yt58* | 45 bp insertion +  5 bp deletion | ACCCGCCCTCTCAAGCTGGTCTACAGTATCTGGCTCTGGATCTACATCTCTCTGGATCTCTGGATTATTACGGGATCTACAGTATCTGGC |
| *rde-3* | *yt59* | 11 bp deletion | ACCCGCCCTCTCAAGCTGGTTATTACGGGATCTACAGTATCTGGCGTGGGC |
| *rde-3* | *yt60* | 3 bp deletion | CCCGCCCTCTCAAGCTGGTTATTACGGGATCTACAGTATCTGGC |
| *rde-3* | *yt61* | 11 bp deletion | TTTCTCACCCGCCCTCTCAAGCTGGTTATTACGGGATCTACAGTATCTGGC |
| *rde-3* | *yt62* | 7 bp deletion | CTCACCCGCCCTCTCAAGCTGGTTATTACGGGATCTACAGTATCTGG |
| *rde-3* | *yt63* | 5 bp deletion | CACCCGCCCTCTCAAGCTGGTTATTACGGGATCTACAGTATCTGG |
| *rde-3* | *yt64* | 1 bp insertion +  11 bp deletion | TCACCCGCCCTCTCAAGCTGTGTTATTACGGGATCTACAGTATCTGGCGTGG |
| *rde-1.1* | *yt53* | 288 bp deletion | ATTGATGAGCCTTTAATTATTATTCATAAACTATTGTAGAGTGAATCTGATGAAAGTTTGGTCAAATTGGAAGTTAAAAAGGAGTCAGGAGATTTCTTCCTCTCCCGAGCGATGGAAACGTCAACAAAGAAAATTCGATTCGTTGTATATAAGCTAACGGATGCGATTGATGAATCGGAAATCAGGTATGCTTTTCAATATGTTTTTTTTATTAATCATCGGTTTTAGCAATTTCAATATTTTCCAGAGAATTCTATCGCCAATTGGTGAAAAAATGCGTGACACGAGGTTTCAACGTGCCCGTAGTAGACCATCAAAACCCTCCTAT |
| *rde-1.1* | *yt54* | 288 bp deletion +  402 bp insertion | ATTGATGAGCCTTTAATTATTATTCATAAACTATTGTAGAGTGAATCTGATGAAAGTTTGGTCAAATTGGAAGTTAAAAAGGAGTCAGGAGATTTCTTCCTCTCCCGAGCGATGGAAACGTCAACAAAGAAAATTCGATTCGTTGTATATAAGCTAACGGATGCGATTGATGAATCGGAAATCAGGTATGCTTTTCAATATGTTTTTTTTATTAATCATCGGTTTTAGCAATTTCAATATTTTCCAGAGAATTCTATCGCCAATTGGTGAAAAAATGCGTGACACGAGGTTTCAACGTGCCCGTAGTAGACCATCAAAACCCTCCTATTTATAAGAAAGAGAGAATTGGAAGATACGACGACATTAAAATTGCACAAATGATGTGCGGCTATGCATCCCGATTAATTCGTTTGAATATTCGTTTATATATTCGAATAATCAGGATGATTTGGCTCTTTTCGATCAATCTGGCAAATCGGATGACGAATTGCTTGTGTTTCTCTTCTTCACGAAAATGGTCGATGAATTATATGGTATGCTGATTTGATCATATTAGATATTTCTCAATATATTGAAGCAACCATTGATCATTGATAGGGCAGATCAAGTATCACTGTGATATTCTTCACGGTGTCGTANTTCCAAGTGGATTAAAATTGCACAAATGATGGCGGTATGCATCCCGATTAATTCGTTTGAATATTCGTTTCAAACGAATTAACGAATTAATTTATAAGAAAGAGAGAATTGGAAGATACGACGACATTAAAATTGCACAAATGATGGCGGTATGCATCCCGATTAA |
| *rde-1.1* | *yt55* | 29 bp insertion | GAGGTTTCAACGTGCCCGTACCATCAAAATAGATGGTACGTCAACGTGCCCATCAAAACCCTCCTATTT |
| *rde-1.1* | *yt56* | 11 bp deletion | ACACGAGGTTTCAACGTGCCCGTAGTAGACCATCAAAACCCTCCTATTTAT |
| *rde-1.1* | *yt57* | 31 bp insertion | AGGTTTCAACGTGCCCGTAGACCATCAAAACCCTCCTATTTATCATCAAAACCATCAAAACCCTCCTATTT |
| *rde-1.2* | *yt65* | 11 bp deletion | AAACTTGTAGACAAGTGTAAATCCCGCGGAATGAGAGTCTCTACACACAAT |
| *rde-1.2* | *yt66* | 10 bp deletion | ACTTGTAGACAAGTGTAAATCCCGCGGAATGAGAGTCTCTACACACAATC |
| *rde-1.2* | *yt67* | 27 bp deletion | ATTTTAGGTCATTCTGCCGCAAACTTGTAGACAAGTGTAAATCCCGCGGAATGAGAGTCTCTACACA |
| *rde-1.3* | *yt68* | 30 bp insertion + 16 bp deletion | TGGCTGAAGTTGAATGTCAATTTTATAAAATCGTAAAATCGTAAAATCGAGCTGGGAGGAGTGAATCAAATTGTCGATTTTACGTA |
| *rde-1.3* | *yt69* | 3 insertions +  14 deletion | CACATTTGGCTGAAGTTGAAGTTTGTCAAGCTGGGAGGAGTGAATCAAATTGTCGAT |
| *rde-1.3* | *yt70* | 22 deletion | CCACATTTGGCTGAAGTTGAATGTCAAGCTGGGAGGAGTGAATCAAATTGTCGATTTTACGT |
